# GeneEmbedLLM: Harnessing Large Language Models for Gene Functional Embedding and Association Prediction

**DOI:** 10.64898/2026.11.05.764758

**Authors:** Ziyan Zhang, Liu Yang, Zhien He, Junxiang Ding, Yixue Li, Rongbo Shen

## Abstract

Effective and high-quality gene function representations are crucial for bioinformatics tasks. While biological knowledge about genes is inherently textual, encompassing functional descriptions and pathway annotations, current methods often rely on single-source, non-textual data, failing to capture the rich semantic information in multi-source gene annotations. In this paper, we propose GeneEmbedLLM, a Large Language Model (LLM)-powered framework for generating universal, semantically-aware gene embeddings. The proposed approach advances the gene representation learning through *multi-source structured text integration* from five authoritative databases and *attribute-aware attention* based on Longformer architecture and contrastive learning. Experimental results on diverse downstream tasks, ranging from molecular interactions and phenotypic dependencies to epigenetic regulation, confirm the validity and effectiveness of our proposed approach, where competitive performance is achieved across all evaluated benchmarks. We further validate the quality of the generated embeddings through comprehensive analyses, including embedding visualizations, intra-pathway similarity assessments, ablation study, and parameter analysis.

## I. Introduction

Characterizing the functional landscape of genes in complex biological systems has become a central challenge in the post-genomic era [1]–[3]. High-quality embedding representations of gene function are crucial for diverse downstream tasks such as gene association prediction and disease gene discovery. In recent years, inspired by advances in Natural Language Processing (NLP), researchers have begun to treat genes as “*semantic entities*”, employing deep learning frameworks to learn low-dimensional dense embeddings from sequences, co-expression relationships, ontology structures, expression profiles, or textual summaries [4]–[6]. However, existing text-based gene embeddings still primarily rely on concise summaries, leaving fine-grained and multi-attribute biomedical annotations underutilized.

To bridge the gap between superficial summaries and comprehensive knowledge, in this paper, we propose GeneEm-bedLLM, a model that leverages the Longformer architecture and contrastive learning to encode gene function with biological text aggregated from multi-source data. Specifically, we first develop a multi-source data integration pipeline with Qwen-Plus. Heterogeneous attributes from five databases are filtered and imputed, resulting in a merged gene attribute table. Unlike summary-only representations such as GenePT, GeneEmbedLLM systematically integrates fine-grained, multi-attribute structured text, including pathway, tissue, cellular-context, Gene Ontology, and clinical prognostic fields. We then use a semi-structured template, designed with special tokens to represent distinct attributes, to transform the tabular data into natural language. Through semantic-preserving augmentation and progressive negative mining, positive and negative samples are generated. Finally, a Longformer [7] encoder equipped with an attribute-aware attention head tailored for biological knowledge learns a gene function representation space via contrastive learning. Experimental results on diverse downstream tasks (i.e., interaction prediction and functional property prediction) confirm the validity and effectiveness of our proposed approach, where competitive performance is achieved on both datasets. To demonstrate that the embeddings obtained from GeneEmbedLLM provide high-quality gene representations capable of reflecting subtle functional distinctions, we further performed embedding visualizations, intra-pathway similarity analysis, robustness and parameter sensitivity analysis. The code and data are available at https://github.com/rheeh/GeneEmbedLLM.

In summary, the contributions of this paper are as follows:

i. We integrate fine-grained gene attributes from multi-source biomedical databases into structured textual descriptions, enabling comprehensive use of specialized knowledge.
ii. We propose GeneEmbedLLM, a transformer-based model that learns gene embeddings via contrastive learning enhanced by semantic-preserving augmentation, progressive negative mining and an attribute-aware attention mechanism tailored for biological attributes.
iii. We demonstrate the model’s effectiveness and superior ability to capture gene function across multiple downstream tasks.

## II. Related Work

Learning high-quality gene embeddings is crucial for downstream tasks such as gene association prediction and disease gene discovery. Early representation methods constructed gene corpora from structured or sequential data. Gene2Vec [8] learned word2vec-like embeddings from gene co-expression patterns, HiG2Vec [9] embedded Gene Ontology (GO) hierarchies in hyperbolic space, and DNABERT [10] modeled DNA sequences through *k*-mer tokenization and transformer pre-training. These approaches capture useful co-expression, ontology, or sequence signals, but they do not directly evaluate how well embeddings understand biomedical textual annotations. Single-cell foundation models have also advanced rapidly. Geneformer and scGPT learn representations from large-scale expression profiles and are effective for dosage, perturbation, and cell-state-related tasks [11]–[13]. Their strength lies in modeling dynamic transcriptional context, whereas our work focuses on static functional semantics encoded in curated biomedical text. These two directions are therefore complementary rather than interchangeable. Recent LLM-based methods have shown that textual gene knowledge can provide useful priors. GenePT [14] encodes gene names and NCBI summaries using text embeddings, demonstrating that even simple summary-based text representations can support single-cell and gene-level analyses. X-Cell further incorporates GenePT embeddings as one of several cross-attended biological priors, together with protein-language, interaction-network, dependency, and morphology-derived priors, for causal perturbation prediction across cellular contexts [15]. However, summary-only descriptions omit many fine-grained attributes distributed across specialized databases, including pathway membership, tissue or cell-type context, molecular function, biological process, and prognostic evidence. GeneEmbedLLM addresses this gap by systematically integrating multi-source, multi-attribute structured text and by using attribute-aware attention to aggregate biologically meaningful fields. Following this scope, our benchmark focuses on methods that are directly informative for evaluating text-derived gene semantics. We compare against two representative LLM-based methods: GenePT (text-embedding-ada-002) and Gemma (a general-purpose LLM). We also include Gene2Vec as a widely used static gene embedding baseline. Other categories, such as sequence-based foundation models (e.g., DNABERT) and GO-based approaches (e.g., HiG2Vec), operate on raw sequences or ontology structures rather than textual annotations, and are therefore not directly comparable for evaluating semantic understanding from biomedical text.

## III. Method

As illustrated in Figure 1, *GeneEmbedLLM* learns gene representations by aligning heterogeneous biological knowledge into a unified semantic space. The pipeline integrates multi-source gene attributes, serializes them into structured text, and encodes them with a parameter-efficient Longformer optimized by contrastive learning.

**Figure 1.**
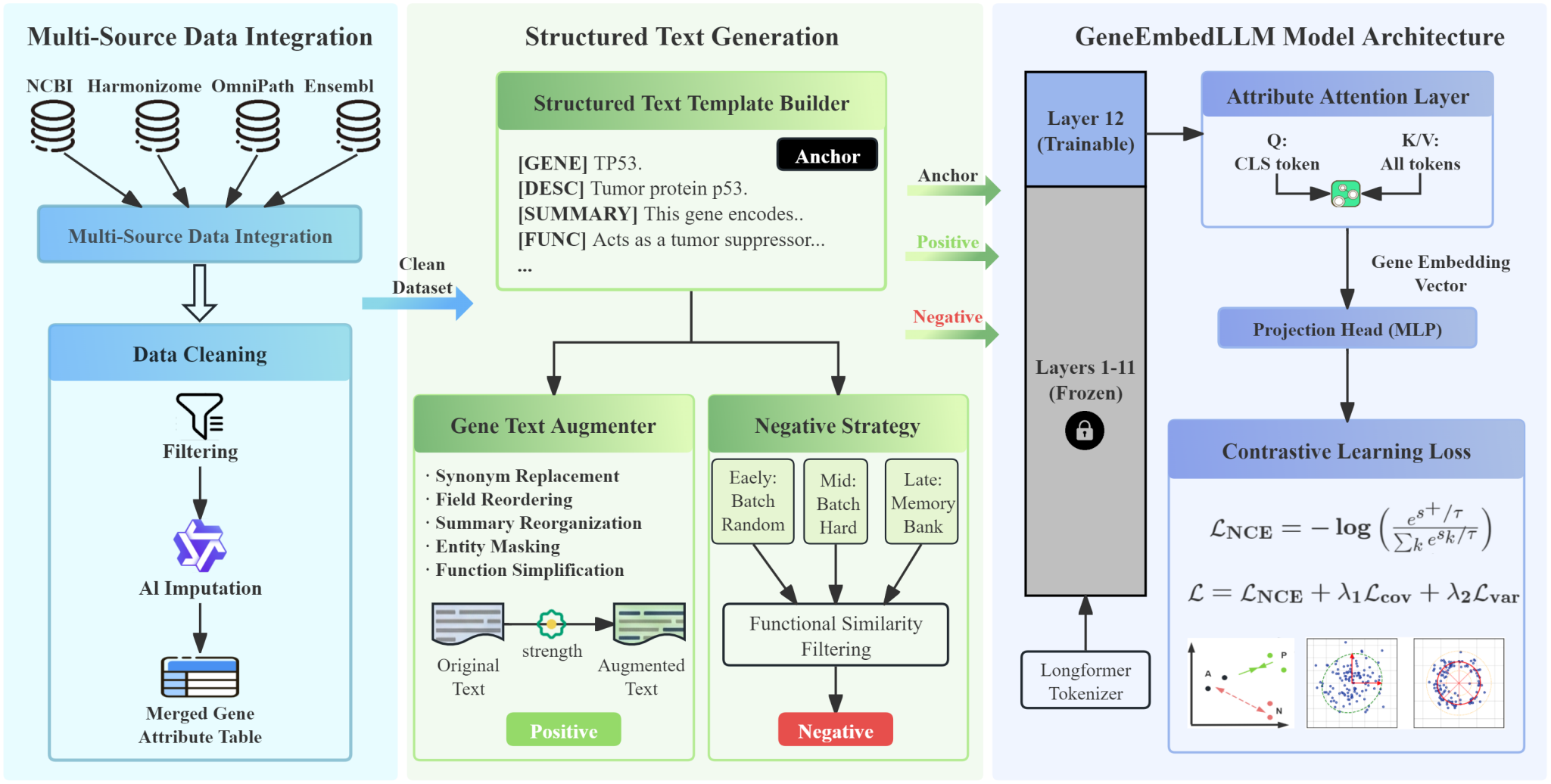
The overall framework of GeneEmbedLLM. The pipeline consists of three cohesive modules: (1) Multi-Source Data Integration, which aggregates, filters, and imputes heterogeneous gene attributes from five authoritative databases using Qwen-Plus; (2) Structured Text Generation, which transforms tabular data into semantically rich text sequences via attribute-aware templates, employing semantic-preserving augmentation and progressive negative mining; (3) GeneEmbedLLM Model Architecture, which maps these sequences into an embedding space using a parameter-efficient Longformer encoder equipped with an attribute-aware attention head and optimized via regularized contrastive learning.

### A. Multi-Source Data Integration

To construct a comprehensive semantic landscape, we aggregated heterogeneous textual data from five authoritative biological databases: *Harmonizome, OmniPath, Ensembl, NCBI Gene, and the Human Protein Atlas*. Preprocessing involved case-insensitive deduplication, non-human pathway filtering, and robust error handling, resulting in 19,773 curated gene records. To mitigate semantic gaps arising from uneven field coverage across nine core attributes, we employed *Qwen-Plus* for targeted missing-entry imputation while preserving originally observed database records. Across these imputed textual attributes, average missingness decreased from 22.7% to 1.7%. To control hallucination risks, we enforced a rigorous *three-tier reliability protocol*: (1) *Constraint-Based Prompting*: Strict system prompts required “well-established” facts from authoritative sources and prohibited speculative content, with temperature set to 0.1 for deterministic output. (2) *Heuristic Uncertainty Filtering*: A blacklist of epistemic uncertainty markers (e.g., “speculative”, “not well characterized”) automatically discarded uncertain generations. (3) *Human Audit*: We cross-checked 180 randomly sampled imputed entries against external literature and assigned *Supported, Partially Supported*, or *Unsupported / Revision Suggested* labels. Among them, 127 entries (70.56%) were supported, 41 (22.78%) were partially supported, and 12 (6.67%) were unsupported or required revision. Overall, 93.34% were at least partially supported, suggesting useful attribute completion while still requiring conservative filtering for uncertain cases. Figure 2 summarizes the field-wise distribution of audit outcomes across imputed attributes.

**Figure 2.**
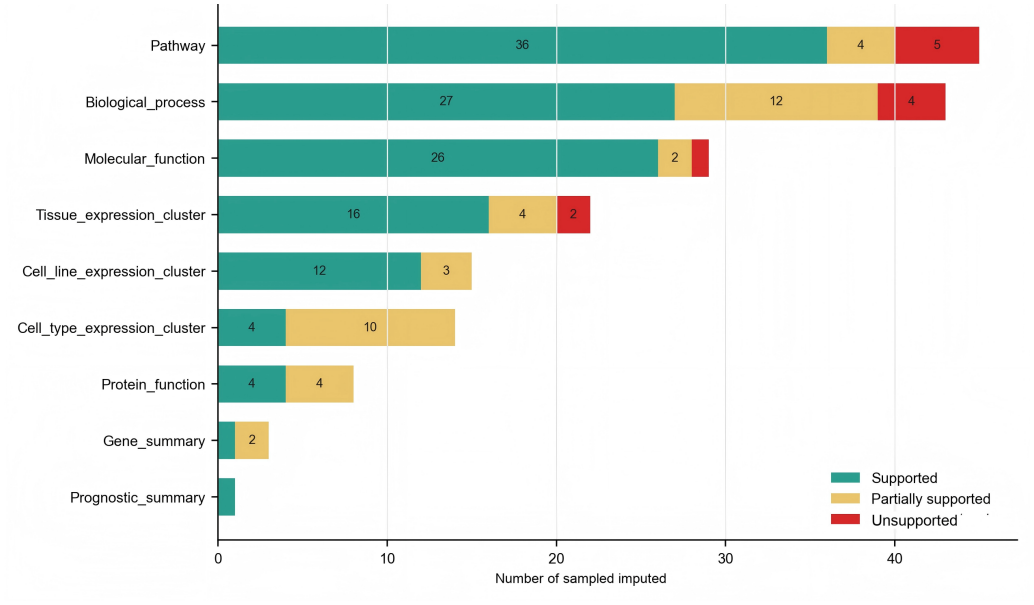
Field-wise manual audit distribution of imputed attributes. The stacked bars show supported, partially supported, and unsupported/revision-suggested entries across audited imputed fields.

### B. Structured Text Generation

a. *Attribute-Aware Template Construction:* We designed a semi-structured serialization template to bridge the gap between tabular data and language models. Distinct attributes are encapsulated within special tokens (e.g., `[GENE]`, `[TISSUE]`, `[PATHWAY]`). This explicit tagging (*S*_raw_ → *S*_struct_) preserves the semantic integrity of independent fields and provides strong structural priors during encoding.
b. *Semantic-Preserving Augmentation:* We introduce *GeneTextAugmenter* to generate semantically consistent positive views through curated synonym substitution, entity masking, non-core field reordering, functional abstraction, and controlled pathway-list variation.
c. *Progressive Negative Mining:* We adopt a curriculum-based negative sampling strategy that evolves dynamically throughout the training process to facilitate robust representation learning:
  - *Phase 1: In-batch Random Sampling (Epochs 0–4)*. This phase establishes foundational feature distributions by randomly selecting negatives within the current batch.
  - *Phase 2: Hard Negative Mining (Epochs 5–7)*. We select negatives with high cosine similarity to the anchor to contrast functionally similar yet distinct genes, thereby sharpening decision boundaries.
  - *Phase 3: Memory Bank Sampling (Epoch 8+)*. We introduce a memory bank (*N* = 4096) to store historical embeddings, decoupling negative sample size from batch limits and enhancing diversity.

Crucially, we incorporate a *False Negative Filtering* mechanism to mitigate semantic collisions. Gene pairs are explicitly excluded from the negative sampling pool if they coexist in the same *KEGG* pathway [16] or exhibit a high interaction score (≥ 700) in *STRING* [17].

### C. GeneEmbedLLM Model Architecture

a. *Parameter-Efficient Long-Context Encoder:* We select *Longformer-base-4096* [7] as the backbone for three reasons: (1) its sliding window attention handles extensive gene profiles with linear complexity; (2) its configurable global attention aligns with our “attribute-aware” design by prioritizing key semantic anchors (e.g., [PATHWAY], [TISSUE]); and (3) its encoder-only architecture is natively optimized for embedding generation. To accommodate these, we expand the vocabulary with 12 domain-specific attribute tokens. To align with biological semantics while preventing overfitting, we freeze the bottom 11 layers and only update the top transformer layer and embedding heads during training.
b. *Attribute-Aware Attention Pooling:* To synthesize a unified gene representation from variable-length, attribute-structured sequences, we replace standard static pooling with a dedicated *Multi-Head Attribute-Aware Attention* mechanism. Let **H** ∈ ℝ^*L×d*^ denote the sequence of hidden states from the top encoder layer. We utilize the global `[CLS]` embedding as a learnable query (**q**_*cls*_), attending to the full sequence, which serves as both key (*K*) and value (*V*). Attribute awareness is induced through domain-specific special tokens (e.g., `[PATHWAY]`, `[TISSUE]`), which serve as semantic anchors and allow different heads to attend to salient biological fields. Formally, the pooling operation is defined as:

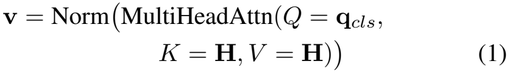

where Norm(·) denotes *L*_2_ normalization. The resulting vector **v** serves as the final gene representation, aggregating heterogeneous attributes in a content-dependent manner.
c. *Projection Head and Multi-Objective Loss:* During the training phase, the normalized gene representation **v** is mapped to a latent contrastive space **z** = MLP(**v**) ∈ ℝ^*d*^ via a non-linear projection head. We optimize the model using a composite objective function ℒ = ℒ_NCE_ + *λ*_1_ ℒ_cov_ + *λ*_2_ ℒ_var_, where each component addresses a specific aspect of the embedding geometry. The primary component, ℒ_NCE_, enforces *alignment and uniformity* by pulling positive pairs together while pushing negatives apart:

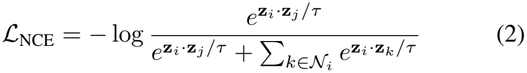

where *τ* is a dynamic temperature parameter. To enhance the information content, we incorporate *feature decorrelation* via ℒ_cov_, which minimizes the squared off-diagonal elements of the batch covariance matrix *C*(**Z**) to prevent dimensional redundancy:

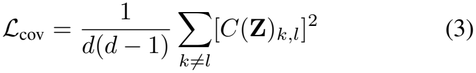

Simultaneously, to mitigate the risk of *representation collapse*, we apply an *anti-collapse constraint* ℒ_var_ using a hinge loss, ensuring the mean variance across dimensions exceeds a target threshold *γ* = 0.05:

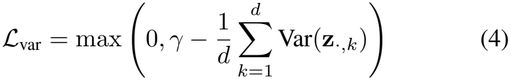

## IV. Experiments

### A. Baselines

Following the comparison scope discussed in Related Work, we benchmark GeneEmbedLLM against three representative embedding models alongside a negative control:

- *Gene2Vec* [8]: A static 200-dimensional embedding model trained on large-scale gene co-expression networks. We utilize the officially released pre-trained embeddings.^1^
- *GenePT* [14]: A text-based model that leverages OpenAI’s `text-embedding-ada-002` to encode gene summaries from NCBI into 1536-dimensional vectors. We use the embeddings from the official v2 release.^2^
- *Gemma* [18]: To evaluate *architectural gains* independent of textual content, we establish a general-purpose baseline using the locally deployed `embedding-gemma-300m`. By feeding the *exact same multi-source gene texts* used in our method into the model, we extract *L*_2_-normalized `[CLS]` vectors as 768-dimensional embeddings.
- *Random Gaussian Embeddings*: We generate dimension-matched (768/1536-dim) vectors from a standard normal distribution. These serve as negative controls to rule out performance gains attributed solely to high dimensionality.

### B. Downstream Tasks and Implementation

To systematically assess embedding quality, we designed benchmarks covering molecular interactions, phenotypic dependencies, and epigenetic regulation.

a. *Interaction Prediction:* We evaluated molecular interaction capabilities using Gene-Gene Interaction (GGI) and Protein-Protein Interaction (PPI) tasks. For GGI, we utilized the benchmark dataset from [8] containing over 200,000 co-expression pairs. For PPI, we employed three datasets constructed by [19]: *HuRI* (52,569 pairs), *Lit-BM* (13,441 pairs), and the comprehensive *HI-union* (64,006 pairs).^3^ To ensure robust evaluation for PPI, we constructed balanced binary classification tasks using *HI-union* as the global positive reference for negative sampling and implemented a *gene-wise split* to prevent data leakage. For both GGI and PPI, the feature representation for a pair (*g*_*i*_, *g*_*j*_) is constructed by summing their individual embeddings: **v**_*pair*_ = **v**_*i*_ + **v**_*j*_.
b. *Functional Property Prediction:* We extended the evaluation to gene-level functional properties. First, we conducted Gene Essentiality Prediction using binary annotations sourced from the Open Targets platform (v25.09) [20].^4^ Second, following [11], we evaluated four Chromatin Dynamics and Dosage Sensitivity tasks: distinguishing bivalent vs. non-methylated genes, Lys4-methylated vs. non-methylated genes, dosage-sensitive vs. insensitive TFs, and long-range vs. short-range TFs.^5^ These tasks assess the model’s capacity to encode deep genomic regulatory information.
c. *Training and Evaluation Details:* GeneEmbedLLM was trained on a single *NVIDIA H800 PCIe* GPU (80GB VRAM). Each epoch required approximately 40 minutes, resulting in a total training time of approximately 16 hours for the complete 24-epoch procedure. We employed mixed-precision training (FP16) to optimize computational efficiency and memory utilization. Although training spanned 24 epochs, we selected the checkpoint from epoch 16 as the final model for all evaluations, as it demonstrated peak performance across validation benchmark tasks. The backbone is *Longformer-base-4096* [7], which contains approximately 149 million total parameters. We added 12 attribute tokens, a 12-head attribute attention layer, and a two-layer projection head. By freezing the bottom 11 transformer layers, trainable parameters constitute only approximately 8–10% of the total model parameters. The main hyperparameter settings are reported in Table I. For all downstream tasks, we evaluated each embedding with both Logistic Regression (LR) and K-Nearest Neighbors (KNN) classifiers. Reporting both classifiers allows us to distinguish representation quality from classifier-specific effects, while keeping the downstream evaluation protocol consistent across embeddings. We report the average performance of 5-fold cross-validation for all tasks, excluding GGI which utilizes a fixed dataset split.

## Results

### A. Visualization of Gene Embeddings

To qualitatively evaluate the topological properties of the learned embedding space, we project the 768-dimensional embeddings of 19,773 human genes into a 2D plane using UMAP (Figure 3), color-coded by 14 functional categories. The visualization reveals a globally organized manifold where genes belonging to the same functional category exhibit discernible spatial aggregation. While major functional groups maintain clear spatial separation within the latent space, we also observe intrinsic overlaps between certain categories. These overlapping regions likely reflect inherent biological pleiotropy and functional interconnectedness, where specific genes participate in multiple overlapping pathways. Overall, this spatial arrangement demonstrates that GeneEmbedLLM effectively maps complex biological semantics into geometric proximity, confirming the model’s ability to capture functional relationships during the unsupervised training process.

**Figure 3.**
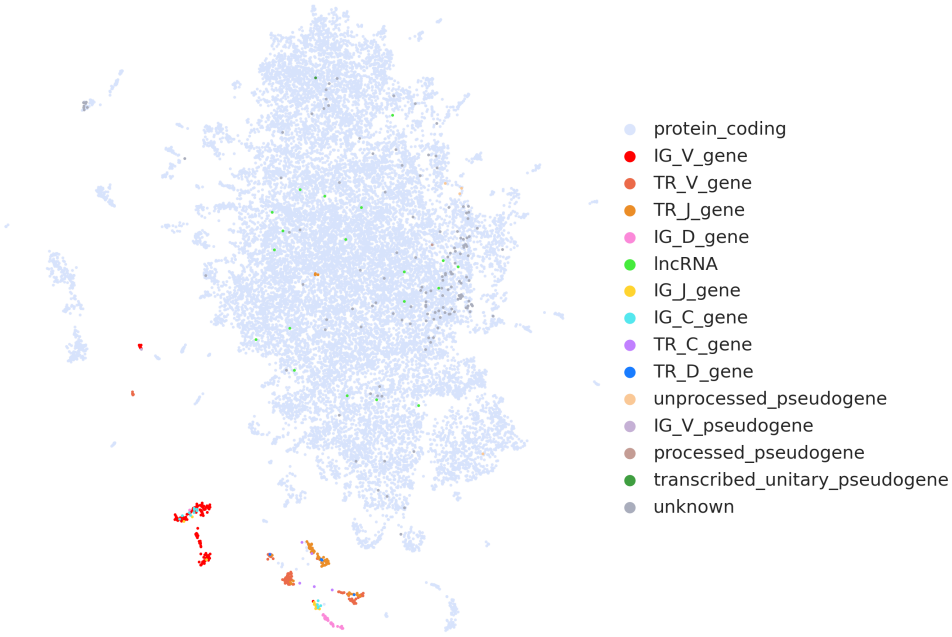
UMAP visualization of gene embeddings colored by functional categories.

### B. Performance on Interaction Prediction

We evaluated GeneEmbedLLM on interaction prediction tasks, including GGI and PPI, to examine whether the learned embeddings capture biologically meaningful functional associations. As summarized in Table II, both LR and KNN results are retained for comparison. GeneEmbedLLM (KNN) achieves the best ROC-AUC on the GGI benchmark and also obtains the strongest performance on the literature-curated PPI (Lit-BM) benchmark. For PPI prediction, GeneEmbedLLM exhibits stable and competitive performance across datasets with distinct construction biases. On the HuRI benchmark, which focuses on experimentally validated interactions, GenePT (KNN) obtains the highest ROC-AUC, while GeneEmbedLLM remains close to the leading text-based baselines and achieves competitive F1-score under both classifiers. This suggests that biologically grounded attributes, such as pathway participation and cellular context, provide complementary signals for identifying true interactions under sparse supervision. The slight advantage of GenePT/Gemma on HuRI may reflect the fact that experimentally validated binary interactions are not always explicitly recorded in pathway- or GO-centered attributes, whereas broad text embeddings can retain more diffuse association cues from summaries. On the literature-curated Lit-BM dataset, GeneEmbedLLM clearly outperforms all compared baselines, achieving the best accuracy, F1-score, and ROC-AUC. These results indicate that explicitly modeling structured biological attributes contributes information beyond coarse semantic similarity alone, particularly for interaction prediction scenarios where functional context is critical.

### C. Performance on Functional Property Prediction

We extended our evaluation to five gene-level functional property prediction tasks covering gene essentiality and genomic regulation. The results are summarized in TABLE III. For Gene Essentiality prediction, the general-purpose LLM embedding model Gemma achieves the best ROC-AUC among the compared embeddings, while GeneEmbedLLM remains highly competitive. This suggests that broad functional centrality signals relevant to gene indispensability are well captured by large-scale text-based representations. The gap also indicates that essentiality may depend on general biological centrality and disease relevance signals that are not fully captured by our manually structured attribute set. In contrast, for Dosage Sensitivity, GeneEmbedLLM (LR) and Gemma (LR) obtain the strongest rounded ROC-AUC scores, indicating that regulatory properties associated with dosage balance can be effectively captured from structured functional attributes without relying on expression-based supervision. Under KNN, GeneEmbedLLM also achieves the best Dosage Sensitivity performance among the evaluated embedding models. For Bivalent vs. Lys4-methylated classification, Gemma (LR) and GeneEmbedLLM (LR) are tied at the rounded ROC-AUC level, while their KNN variants remain close to the leading text-based models. For Bivalent vs. Non-methylated and TF Range tasks, the fine-tuned expression-based Geneformer achieves the best results among all listed methods, highlighting the advantage of transcriptomic context in modeling chromatin accessibility and transcription factor regulatory range. Among text-based embeddings, GeneEmbedLLM achieves the best ROC-AUC on Bivalent vs. Non-methylated classification, whereas TF Range remains more challenging for static text-derived representations. This result suggests that static knowledge-driven embeddings and dynamic transcriptomic representations capture complementary biological signals, motivating future joint embedding designs. Thus, non-leading results on these regulatory tasks should be interpreted as a boundary of static annotation-derived embeddings rather than a failure of semantic integration.

**TABLE I.** Hyperparameter configuration for GeneEmbedLLM training.

| *Hyperparameter* | *Value* |
|---|---|
| Maximum Sequence Length | 3072 tokens |
| Batch Size / Gradient Accumulation Steps | 10 / 2 |
| Effective Batch Size | 20 |
| Peak Learning Rate | 1e-5 |
| Weight Decay | 1e-4 |
| Total Training Epochs | 24 |
| Temp. Schedule (τ) | max(0.02, 0.4×0.95<sup>epoch</sup>) |
| Cov. Loss Weight (λ<sub>1</sub>) | 0.001 |
| Var. Loss Weight (λ<sub>2</sub>) | 0.0005 |
| Memory Bank Size | 4096 |

**TABLE II.** Performance comparison on interaction prediction tasks. Metrics are reported as mean *±*s.d. (GGI uses a fixed split, thus no s.d. is reported). Best results are bolded.

| Task | Model | Accuracy | F1-score | ROC-AUC |
|---|---|---|---|---|
| **GGI** | Random (768)+LR | 0.49 | 0.41 | 0.49 |
|  | Random (768)+KNN | 0.49 | 0.42 | 0.50 |
|  | Random (1536)+LR | 0.52 | 0.43 | 0.53 |
|  | Random (1536)+KNN | 0.49 | 0.40 | 0.49 |
|  | Gene2Vec + LR | 0.62 | 0.65 | 0.67 |
|  | Gene2Vec + KNN | 0.58 | 0.61 | 0.62 |
|  | GenePT + LR | 0.71 | 0.70 | 0.79 |
|  | GenePT + KNN | 0.73 | 0.74 | 0.80 |
|  | Gemma + LR | 0.72 | 0.72 | 0.79 |
|  | Gemma + KNN | 0.75 | **0.78** | 0.83 |
|  | GeneEmbedLLM + LR | 0.75 | 0.76 | 0.82 |
|  | GeneEmbedLLM + KNN | **0.77** | **0.78** | **0.84** |
| **PPI (HuRI)** | Random (768)+LR | 0.48 ± 0.01 | 0.38 ± 0.01 | 0.47 ± 0.01 |
|  | Random (768)+KNN | 0.50 ± 0.01 | 0.42 ± 0.01 | 0.50 ± 0.01 |
|  | Random (1536)+LR | 0.49 ± 0.01 | 0.37 ± 0.02 | 0.49 ± 0.01 |
|  | Random (1536)+KNN | 0.51 ± 0.00 | 0.42 ± 0.01 | 0.51 ± 0.01 |
|  | Gene2Vec + LR | 0.52 ± 0.02 | 0.47 ± 0.03 | 0.53 ± 0.03 |
|  | Gene2Vec + KNN | 0.53 ± 0.01 | 0.46 ± 0.02 | 0.54 ± 0.01 |
|  | GenePT + LR | 0.59 ± 0.01 | 0.54 ± 0.03 | 0.63 ± 0.02 |
|  | GenePT + KNN | **0.61 ± 0.01** | 0.57 ± 0.02 | 0.65 ± 0.02 |
|  | Gemma + LR | 0.60 ± 0.01 | 0.55 ± 0.01 | 0.64 ± 0.01 |
|  | Gemma + KNN | 0.60 ± 0.01 | **0.59 ± 0.01** | **0.65 ± 0.01** |
|  | GeneEmbedLLM + LR | 0.60 ± 0.01 | 0.57 ± 0.02 | 0.65 ± 0.02 |
|  | GeneEmbedLLM + KNN | 0.60 ± 0.02 | 0.57 ± 0.02 | 0.65 ± 0.02 |
| **PPI (Lit-BM)** | Random (768)+LR | 0.49 ± 0.01 | 0.40 ± 0.02 | 0.49 ± 0.02 |
|  | Random (768)+KNN | 0.51 ± 0.01 | 0.43 ± 0.02 | 0.51 ± 0.02 |
|  | Random (1536)+LR | 0.50 ± 0.01 | 0.41 ± 0.02 | 0.50 ± 0.01 |
|  | Random (1536)+KNN | 0.50 ± 0.02 | 0.42 ± 0.03 | 0.50 ± 0.03 |
|  | Gene2Vec + LR | 0.58 ± 0.02 | 0.55 ± 0.04 | 0.60 ± 0.03 |
|  | Gene2Vec + KNN | 0.54 ± 0.02 | 0.54 ± 0.02 | 0.55 ± 0.02 |
|  | GenePT + LR | 0.63 ± 0.01 | 0.59 ± 0.02 | 0.67 ± 0.02 |
|  | GenePT + KNN | 0.65 ± 0.03 | 0.63 ± 0.03 | 0.69 ± 0.04 |
|  | Gemma + LR | 0.66 ± 0.01 | 0.63 ± 0.01 | 0.71 ± 0.01 |
|  | Gemma + KNN | 0.61 ± 0.02 | 0.67 ± 0.03 | 0.69 ± 0.02 |
|  | GeneEmbedLLM + LR | 0.64 ± 0.02 | 0.62 ± 0.03 | 0.70 ± 0.02 |
|  | GeneEmbedLLM + KNN | **0.71 ± 0.02** | **0.70 ± 0.03** | **0.77 ± 0.02** |

**TABLE III.** ROC-AUC comparison on gene functional property prediction tasks (mean *±* S.D.). Best results are bolded. Models marked with *†* cite results directly from [11].

| **Model** | **Dosage Sensitivity** | **Bivalent vs. Non-methylated** | **Bivalent vs. Lys4-methylated** | **TF Range** | **Gene Essentiality** |
|---|---|---|---|---|---|
| Random (768)+LR | 0.58 ± 0.08 | 0.57 ± 0.10 | 0.69 ± 0.05 | 0.41 ± 0.11 | 0.50 ± 0.01 |
| Random (768)+KNN | 0.50 ± 0.05 | 0.58 ± 0.06 | 0.59 ± 0.06 | 0.43 ± 0.08 | 0.50 ± 0.01 |
| Random (1536)+LR | 0.56 ± 0.04 | 0.53 ± 0.08 | 0.44 ± 0.09 | 0.54 ± 0.10 | 0.50 ± 0.01 |
| Random (1536)+KNN | 0.46 ± 0.08 | 0.41 ± 0.07 | 0.52 ± 0.05 | 0.52 ± 0.08 | 0.51 ± 0.01 |
| Gene2Vec + LR | 0.89 ± 0.01 | 0.70 ± 0.13 | 0.90 ± 0.03 | 0.62 ± 0.13 | 0.91 ± 0.01 |
| Gene2Vec + KNN | 0.87 ± 0.06 | 0.64 ± 0.15 | 0.89 ± 0.06 | 0.62 ± 0.09 | 0.85 ± 0.01 |
| GenePT + LR | 0.88 ± 0.03 | 0.90 ± 0.03 | 0.93 ± 0.06 | 0.69 ± 0.09 | 0.88 ± 0.01 |
| GenePT + KNN | 0.88 ± 0.04 | 0.84 ± 0.11 | 0.92 ± 0.03 | 0.56 ± 0.17 | 0.86 ± 0.01 |
| Gemma + LR | 0.92 ± 0.02 | 0.91 ± 0.04 | 0.94 ± 0.06 | 0.64 ± 0.12 | **0.93** ± **0.01** |
| Gemma + KNN | 0.90 ± 0.01 | 0.85 ± 0.07 | 0.92 ± 0.04 | 0.61 ± 0.08 | 0.90 ± 0.02 |
| Geneformer(fine-tuned)<sup>†</sup> | 0.91 ± 0.02 | **0.93** ± **0.07** | 0.88 ± 0.09 | **0.74** ± **0.08** | – |
| GeneEmbedLLM + LR | **0.92** ± **0.01** | 0.91 ± 0.05 | **0.94** ± **0.05** | 0.62 ± 0.11 | 0.91 ± 0.01 |
| GeneEmbedLLM + KNN | 0.91 ± 0.02 | 0.87 ± 0.06 | 0.92 ± 0.05 | 0.55 ± 0.11 | 0.89 ± 0.01 |

**TABLE IV.** Ablation and robustness analysis on the GGI task using the LR downstream classifier. We evaluate the impact of knowledge imputation, false negative filtering, and training configurations on interaction prediction performance.

| **Configuration** | **Accuracy** | **F1-score** | **ROC-AUC** |
|---|---|---|---|
| **GeneEmbedLLM (Default)** | **0.748** | **0.762** | **0.821** |
| *Data Construction* |  |  |  |
| w/o Missing Attribute Imputation | 0.744 | 0.754 | 0.815 |
| *False Negative Filtering* |  |  |  |
| w/o False Negative Filtering | 0.743 | 0.755 | 0.817 |
| *Training Strategy* |  |  |  |
| Full Fine-tuning (w/o Frozen Layers) | 0.731 | 0.742 | 0.804 |
| Static Sampling (Random Only) | 0.738 | 0.748 | 0.810 |
| High Regularization (λ × 10) | 0.746 | 0.758 | 0.817 |

### D. Intra-Pathway Similarity Analysis

To evaluate the model’s capability in capturing biological functional relationships, we analyze the embedding topology of the *Glycolysis* pathway. Specifically, we curate a set of *67* genes associated with the Glycolysis pathway from the KEGG database, alongside a control group of *60* randomly sampled genes from the human genome. Figure 4A and B characterize the resulting embedding space generated by GeneEmbedLLM. The heatmap (Figure 4A) reveals a prominent block-diagonal structure: the 67 pathway genes form a dense high-similarity cluster (red block), which is clearly demarcated from the 60 random background genes. This geometric cohesion is further corroborated by the UMAP projection (Figure 4B), where Glycolysis genes (red points) aggregate into a tight, isolated cluster, while random genes (green points) exhibit a diffuse, scattered distribution. To quantify this topological quality, we compare the similarity distributions across different models in Figure 4C. GeneEmbedLLM achieves the most distinct separation between functional signals and background noise. As shown in the boxplots, the median intra-pathway similarity (red, ≈ 0.80) is significantly higher than the random baseline (blue, ≈ 0.5), yielding a substantial discriminative margin of Δ ≈ 0.3. In contrast, baseline models exhibit limited functional resolution. While GenePT produces high intra-pathway scores, it suffers from anisotropy; its signal distribution overlaps with the random background (Δ ≈ 0.03), hindering effective gene discrimination. Gemma shows better separation than GenePT but a weaker overall signal (mean ≈ 0.49). Gene2Vec fails to capture meaningful functional semantics, with almost no margin between groups (Δ ≈ 0.04). These results demonstrate that GeneEmbedLLM uniquely preserves both high intra-pathway cohesiveness and robust separability.

**Figure 4.**
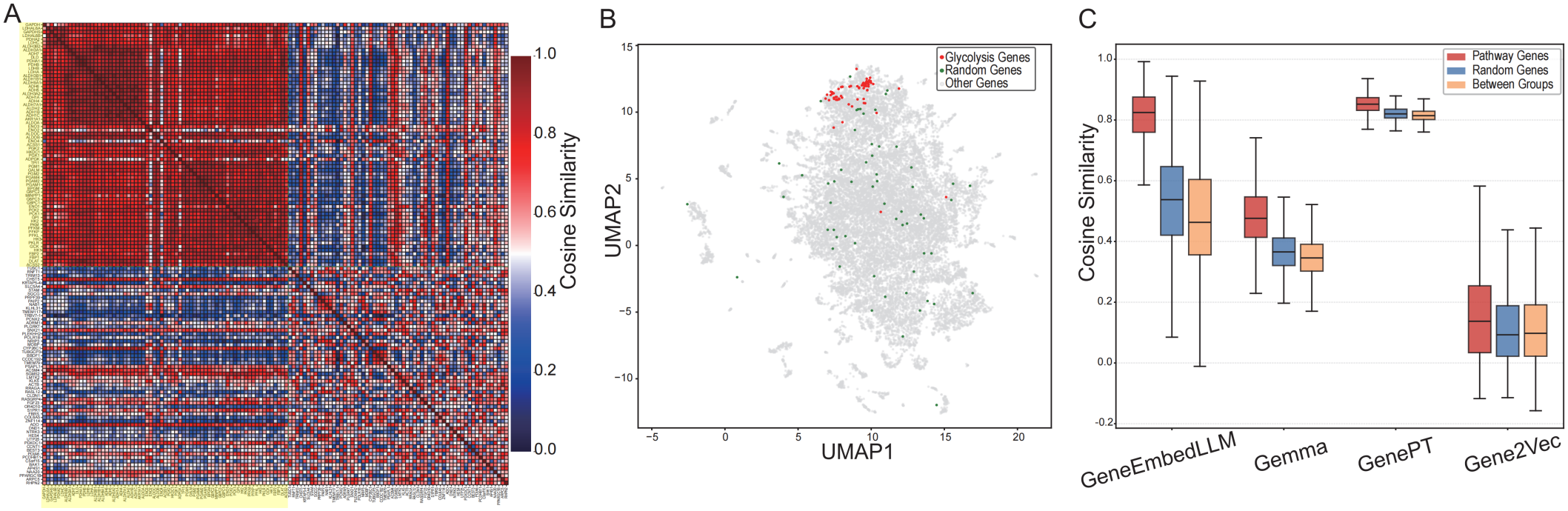
Analysis of intra-pathway functional coherence. **(A)** Cosine similarity heatmap of GeneEmbedLLM for 67 Glycolysis genes (derived from KEGG) and 60 random genes. Yellow-highlighted gene labels on the axes denote intra-pathway genes. **(B)** UMAP visualization showing the spatial aggregation of the 67 Glycolysis genes (red) against 60 random genes (green). **(C)** Comparison of cosine similarity distributions across models, illustrating the discriminative margin (Δ) between functional and random gene sets. In the boxplots, the central line indicates the median, and whiskers extend to the most extreme data points within 1.5*×* the interquartile range (IQR); outliers are omitted for visual clarity.

### E. Ablation and Robustness Analysis

To further assess the contribution of the major design components in GeneEmbedLLM, we conducted ablation and robustness analyses on the GGI task using the LR down-stream classifier by systematically modifying textual input construction, false-negative filtering, training configurations, and regularization settings (Table IV).

a. *Impact of Text Construction:* Removing the missing-attribute imputation module consistently degraded model performance, indicating that targeted knowledge completion improves the semantic completeness of structured gene profiles and provides complementary biological signals.
b. *Impact of False Negative Filtering:* We further evaluated whether the GGI result depends primarily on biologically informed false negative filtering. Removing this filtering slightly reduced ROC-AUC from 0.821 to 0.817 using the same LR downstream classifier, but the resulting model still outperformed the LR variants of Gene2Vec, GenePT, and Gemma on the GGI benchmark. This suggests that the strong GGI performance is not solely attributable to pseudo-negative filtering or potential overlap with STRING/KEGG-derived signals, while the small drop indicates that such filtering still helps reduce semantic collisions during contrastive training.
c. *Impact of Training Strategy:* Alternative optimization strategies also reduced overall performance. Full fine-tuning without frozen lower layers showed weaker generalization, suggesting that parameter-efficient tuning better preserves useful pre-trained semantic priors while reducing overfitting risk. Similarly, replacing progressive negative mining with static random sampling degraded performance, highlighting the importance of curriculum-based hard negative exposure for learning discriminative representations.
d. *Hyperparameter Sensitivity:* Increasing the regularization strength resulted in only marginal performance changes, demonstrating that GeneEmbedLLM remains stable under substantial hyperparameter perturbations. This suggests that the default setting provides a robust balance between representation capacity and generalization.

## VI. Conclusion

We propose *GeneEmbedLLM*, which advances gene representation learning through multi-source structured text integration and attribute-aware attention. Distinct from summary-based approaches such as GenePT, our framework encodes fine-grained biological attributes from curated databases and learns a unified embedding space through contrastive learning. Evaluations across interaction prediction, functional property prediction, and pathway coherence analyses demonstrate that GeneEmbedLLM captures biologically meaningful functional relationships. This study also has limitations. The Qwen-Plus imputation step can improve attribute completeness, but it may introduce hallucinated or outdated facts. In our manual audit, 6.67% of sampled imputed entries were unsupported or required revision; at the 19,773-gene scale, this rate would correspond to roughly 1,300 potentially erroneous entries, mainly concentrated in pathway- and GO-related fields. GeneEmbedLLM is also constrained by the quality, coverage, and update frequency of external databases, and static textual priors may miss dynamic cell-state-dependent regulatory information. In addition, because STRING and KEGG contain interaction or pathway evidence that may overlap with the GGI benchmark, false negative filtering may introduce residual information leakage. The w/o-filtering ablation reduces this concern but does not fully eliminate it, as its performance is slightly lower than the default GeneEmbedLLM setting. Future work will apply GeneEmbedLLM to two downstream biological discovery settings: identifying latent marker genes in cellular contexts and constructing disease-function semantic spaces to support candidate disease-gene insight. We also plan to explore integration with scRNA-seq latent representations and interaction graphs, and to evaluate cross-species generalization while clearly separating human-specific annotations from conserved functional signals.

## Footnotes

1 https://github.com/jingcheng-du/Gene2vec

2 https://zenodo.org/records/10833191

3 https://interactome-atlas.org/download

4 https://platform.opentargets.org/downloads

5 https://huggingface.co/datasets/ctheodoris/Genecorpus-30M

